# Procollagen C-Proteinase Enhancer 1 (PCPE-1) is a marker of myocardial fibrosis and impaired cardiac function in a murine model of pressure overload

**DOI:** 10.1101/2021.03.05.434071

**Authors:** Priscillia Lagoutte, Alexandra Oudot, Mélissa Dussoyer, Victor Goncalves, Mélanie Guillemin, Olivier Bouchot, David Vandroux, Pierre-Simon Bellaye, Catherine Moali, Sandrine Vadon-Le Goff

**Affiliations:** Univ Lyon, CNRS, Tissue Biology and Therapeutic Engineering Laboratory, LBTI, UMR5305, F-69367, Lyon, France; Georges-François LECLERC Cancer Center – Unicancer, Dijon, France; Institut de Chimie Moléculaire de l’Université de Bourgogne, UMR6302, CNRS, Université Bourgogne Franche-Comté, F-21000 Dijon, France; CHU, Dijon, France; NVH Medicinal, Dijon, France

**Keywords:** Cardiac fibrosis, Procollagen C-proteinase enhancer 1 (PCPE-1), PET-MR imaging, circulating biomarker, collagen biosynthesis, heart failure

## Abstract

**Aims:** Procollagen C-proteinase enhancer 1 (PCPE-1) is an extracellular matrix protein and a major regulator of fibrillar collagen biosynthesis. Previous work has shown that its abundance is often increased in the context of tissue repair and fibrosis. The present study was designed to evaluate its potential as a biomarker of myocardial interstitial fibrosis (MIF), a well-established pathogenic pathway leading to heart failure.

**Methods and Results:** Cardiac fibrosis was induced in rats using an optimized model of chronic pressure overload triggered by angiotensin II and N^ω^-nitro-L-arginine methyl ester (L-NAME). All treated animals suffered from heart hypertrophy and the increase in heart collagen volume fraction (CVF), evidenced by histology and ^68^Ga-Collagelin uptake, confirmed the development of cardiac fibrosis. Functional analysis by simultaneous PET-MRI further showed that our model closely reflected the pathological features seen in human MIF, including left ventricle thickening and diastolic dysfunction associated with decreased ejection fraction. PCPE-1 mRNA and protein levels were augmented by factors of 3.4 and 6.1 respectively in the heart tissue of treated rats. Moreover, protein abundance was well-correlated with CVF (r=0.92, p<0.0001) and PCPE-1 immuno-detection mainly localized the protein to fibrotic areas. Finally, PCPE-1 plasma levels measured by ELISA were increased in fibrotic rats compared to controls.

**Conclusion:** Together, our findings demonstrate that PCPE-1 levels in the heart and circulation tightly reflect the cardiac fibrosis status and heart function impairment in rats and suggest that it could be a very useful marker to monitor human heart diseases leading to fibrosis.

## (1) Introduction

Heart failure (HF) is a major cause of morbidity and mortality worldwide and the most frequent source of hospitalization for patients over 65 years of age^1^. Myocardial interstitial fibrosis (MIF) is one of the main pathogenic factors that predispose to HF^2,3^. Resulting from a variety of acute or chronic injuries, it is characterized by an aberrant and persistent tissue remodeling response, leading to an excessive accumulation of extracellular matrix (ECM) produced by activated fibroblasts (myofibroblasts)^4,5^. As opposed to the large scars resulting from myocardial infarction, interstitial fibrosis is characterized by diffuse and patchy areas located around cardiomyocytes or vessels^6^. This extensive tissue remodeling impairs normal physiological heart function, and results in myocardial stiffening, mechanical, electrical and/or vasomotor dysfunctions that can ultimately lead to death^2,7–9^.

Regardless of the etiology of the disease, fibrillar collagens I and III are the major components of the interstitial ECM observed in the fibrotic myocardium^5,10^. Although it has been recognized for a while that preventing their accumulation could be a valuable strategy to prevent or treat fibrosis^11^, there is presently no therapeutic candidate capable of blocking the critical steps of the collagen biosynthesis pathway. Also, while numerous biomarkers of myocardial fibrosis have been investigated, very few of them have been clinically validated. Hence, there is still an urgent need for new strategies, such as the use of circulating biomarkers or non-invasive imaging methods, that could accurately define the cardiac fibrosis status in patients and provide early and sensitive diagnosis^7,8,12–14^.

Collagen deposition requires proteolytic maturation of the fibrillar collagen precursor molecules also known as the procollagens. This step is carried out by procollagen N- and C-proteinases in the extracellular environment^15^. The bone morphogenetic protein-1 (BMP-1)/tolloid-like proteinases (BTPs) are the main proteases responsible for the removal of the C-propeptides of fibrillar procollagens I-III, the rate-limiting step in collagen fibril formation^16^. They are assisted in this function by a potent and specific enhancer of procollagen C-terminal maturation called procollagen C-proteinase enhancer 1 (PCPE-1)^17–19^.

PCPE-1 is a secreted glycoprotein encoded by the *PCOLCE* gene in humans. It is composed of two CUB (Complement-Uegf-BMP-1) domains and one C-terminal NTR (Netrin-like) domain. The latter can bind to heparin, heparan sulfate proteoglycans and fibronectin^20,21^, while the CUB1CUB2 region is necessary and sufficient for PCPE-1 enhancing activity. Indeed, PCPE-1 can increase the catalytic efficiency of BMP-1 on procollagens more than 10 fold^22,23^ while having no effect on the cleavage of other BTP substrates^18^. This strong collagen specificity can be explained by the ability of PCPE-1 to bind the C-terminal domain of procollagens (C-propeptide) which is later released upon proteolytic maturation. This tight interaction is the basis for the enhancing activity of PCPE-1 as it allows the local unravelling of the procollagen trimer which is thought to facilitate the C-propeptide cleavage by the BTPs^22,24^.

The link between PCPE-1 up-regulation and fibrosis has already been demonstrated in several organs such as liver, muscle, skin or cornea^25–28^. Regarding cardiac fibrosis, PCPE-1 was found to be increased in endomyocardial biopsies from a cohort of 28 patients suffering from aortic valve stenosis^29^. Also, Kessler-Icekson and co-workers used a rat model of myocardial infarction induced by coronary artery ligation to show that *Pcolce* RNA and derived protein were both significantly elevated in the fibrotic heart^30^. They also observed a good correlation between the PCPE-1 and collagen I protein levels, both quantified from immunoblots, and showed that the aldosterone receptor antagonist spironolactone could reverse PCPE-1 up-regulation in this model. Since then, several other studies have reported PCPE-1 up-regulation in myocardial remodeling induced by various stimuli^31–33^. However, the thorough analysis of the ability of PCPE-1 levels to reflect the extent of fibrosis and the consequences for cardiac function is presently missing in a model of MIF.

In this study, we have evaluated the capacity of PCPE-1 to be used as a biomarker of interstitial fibrosis. For this purpose, we have adapted a rat model of cardiac fibrosis using pharmacologically-induced chronic pressure overload. The combined and extensive analysis of heart function, fibrosis and PCPE-1 expression and localisation led to the conclusion that PCPE-1 levels in the heart and circulation tightly reflect cardiac fibrosis and heart function impairment.

## (2) Methods

### Cardiac fibrosis rat model

All animal studies were conducted in accordance with the legislation on the use of laboratory animals (directive 2010/63/EU) and were approved by an accredited ethical committee (C2ea Grand Campus n°105) and the French ministry of research (authorization #2349).

The rat model of cardiac fibrosis used in the present study was induced by pressure overload with minor modifications of a previously described protocol^34^. After 1-week acclimation, Male Wistar Han male rats (250-275 g, Charles River-France) from the “AngII + L-NAME” group (n=8) were given N^ω^-nitro-L-arginine methyl ester (L-NAME, Sigma-Aldrich) in drinking water (40 mg/kg/d) for 4 weeks. After two weeks, the animals were anesthetized under isoflurane (2-2.5 %) and osmotic minipumps (Alzet model 2002, Charles River-France) loaded with angiotensin II (AngII, Sigma-Aldrich) were placed subcutaneously in the interscapular region to achieve 175 μg/kg/d AngII administration during the last 2 weeks of the procedure. Control group animals (n=6) received normal drinking water and sham operation.

The rats were randomized into each experimental group according to their body weight but pairs of rats living in the same cage were not separated. During the 4 weeks of the treatment, one rat from the “AngII + L-NAME” group was euthanized because it reached ethical endpoints (15% body weight loss between day 14 and day 19). Thus, the results of the present study correspond to n=7 for the “AngII+L-NAME” group and n=6 for the control group.

### PET-MR Imaging

Simultaneous PET-MR imaging was performed after 4 weeks of treatment on a fully integrated system (MR Solutions, Guildford, UK) consisting of a 7 T dry magnet (Powerscan MRS-7024-PW) coupled with a SiPM-based dual ring PET system. Animals from each experimental group were distributed all along the imaging experiment duration to reduce the potential bias due to imaging time. A collagelin analog (CPGRVMHGLHLGDDEGPC) was synthesized, bioconjugated with NODAGA and radiolabelled with ^68^Ga, as previously described^35^. PET acquisitions were performed in the list mode from 30 to 60 min after injection of 5 nmol of ^68^Ga-radiolabeled collagelin (20.6 ± 0.8 MBq). MRI cardiac cine acquisitions were simultaneously performed with PET. Images were respiratory- and cardiac-gated using a pneumatic cushion, subcutaneous needles and dedicated software (PC Sam, SAII, Stony Brook, US). Twelve temporal frames per cardiac cycle were acquired. A spoiled gradient recoiled echo - fast low angle shot (FLASH GRE) cine sequence was used with a repetition time (TR) of 10 ms, an echo time (TE) of 3 ms, a flip angle of 40°, a matrix size of 256 × 256 pixels, a voxel size of 0.23 × 0.23 × 1.5 mm^3^ and 4 signal averages. MR left ventricular volumes and thicknesses were determined by manually delineating the endocardial and epicardial contours on short axis slices (AnimHeart, CASIS, Dijon, France) on end-diastolic and end-systolic frames to determine respectively end-diastolic volume (EDV) and end-systolic volume (ESV). Left ventricle volume and thickness were measured in end-diastolic MR images. Left ventricular ejection fraction was calculated from EDV and ESV. The investigator who performed cardiac function analyses was unaware of the treatment (blinded files).

### Blood sampling

At the time of euthanasia (4 weeks), blood was harvested from the tail vein in citrate-coated tubes, centrifuged and plasma samples were frozen at −80°C.

### Cardiac tissue processing – Fibrosis evaluation

At the end of imaging, the rats were euthanized by an intraperitoneal injection of 140 mg/kg pentobarbital (Euthasol Vet^®^, Dechra Veterinary products) and the heart tissue was collected and weighed after atrial tissue removal. The ventricular tissue was cut into 3 parts: the central part was weighed and placed in 10 % neutral-buffered formalin and the basal and apical thirds of the ventricle were flash-frozen and kept at −80°C. Cardiac ^68^Ga-Collagelin uptake was measured with a scintillation *γ*-counter (Cobra 4180, PerkinElmer, Waltham, MA) in the central third of the heart. Results were corrected for heart weight, injected dose of ^68^Ga-Collagelin and radioactive decay and were expressed as a percentage of injected dose (% ID).

After 48-72 h neutral-buffered formalin fixation, heart tissue was paraffin-embedded for histological fibrosis evaluation by Masson trichrome and picrosirius red staining. Left ventricular fibrosis was determined using the ImageJ software v 1.48 by thresholding the specific staining after removal of vascular structures and expressed as percentage of tissue area. Four fields of view per section (magnification x5) were analysed for fibrosis staining and averaged to obtain a mean value per rat and per treatment condition. The investigator who performed fibrosis quantification was unaware of the treatment.

### RNA extraction from tissues and qRT-PCR

Frozen rat heart samples were resuspended and lysed in Nucleozol buffer (740404; Macherey Nagel). Protein were then precipitated with nuclease-free water. Samples were centrifuged (15,000 g; 30 min; 4°C) and supernatants were collected. Total RNA was then purified using the NucleoSpin RNA set of the Nucleozol kit (740406; Macherey Nagel) according to the manufacturer’s recommendations. To eliminate all residual DNA contaminants, DNAse treatment was carried out using the Turbo DNA free kit (AM1907; Ambion). RNA was quantified using a nanodrop ND-2000 spectrophotometer (Thermofisher). RNA quality and integrity were checked by electrophoresis on 1.5 % agarose gels. Purified RNA was aliquoted and stored at −80°C.

200 ng of mRNA were reverse-transcribed to cDNA using the PrimeScript^TM^ RT reagent kit (RR037A; Takara). Fluorescence-based quantitative real-time PCR (qRT-PCR) was then performed using the FastStart Universal SYBR Green Master mix (4913850001; Roche) and a Rotor-Gene Q instrument (Invitrogen). The PCR thermocycler conditions included an initial denaturation (95°C for 10 min) followed by 50 amplification cycles, each comprising denaturation (95°C for 10 s) and annealing (60°C for 20 s) steps. Fluorescence was measured at the end of the annealing step of each cycle and quantification cycles (Cq) were determined for each gene and condition. At the end of the amplification, a melting curve was recorded by heating slowly at 0.5°C/s from 70°C to 94°C. All primers were obtained from Sigma (Table 1). All experiments were performed in three technical replicates. Relative quantification was performed using the 2^−ΔΔCq^ method, which allows the comparison of two conditions after normalization with the reference gene *Rpl32* (Ribosomal protein L32)^36^.

**Table 1.**
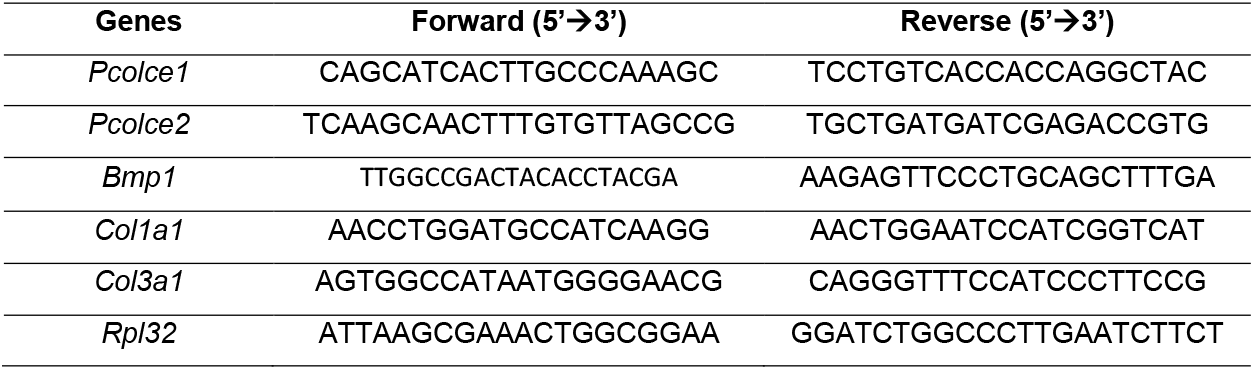
qRT-PCR oligonucleotides.

### Protein extraction from tissues and immunoblotting

Frozen rat heart samples (around 100 mg each) were mechanically broken and resuspended in Laemmli’s sample buffer (0.325 M Tris pH 6.8; 2.5 % SDS; 5 % Glycerol) supplemented with 4.5 % β-mercaptoethanol and 0.5 % bromophenol blue. Proteins were then solubilized by heating for 10 min at 100°C. The heat-denatured samples were rotated for 30 min at room temperature before centrifugation (15,000 g, 15 min). The supernatants were collected and stored at −80°C.

Protein were separated by SDS-PAGE (4-20% acrylamide gradient; Mini-Protean TGX Stain-free gels; Biorad) for the detection of PCPE-1. Equal volumes of protein extracts were loaded in each lane. After migration, gels were activated by UV exposure for 1 minute using a FX Fusion CCD camera (Vilber Lourmat). Proteins were then electrotransferred to PVDF membranes using 10 mM CAPS buffer (pH 11) supplemented with 10 % ethanol. Total protein amount was estimated by fluorescent detection of tryptophan residues using the trihalo compound included in the gel and transferred onto the membrane after activation (stain-free technology). Membranes were blocked for 2 h at room temperature with 5 % skimmed milk in PBS buffer (137 mM NaCl; 2.7 mM KCl; 10 mM Na_2_HPO_4_; 1.8 mM KH_2_PO_4_ pH 7.4). Membranes were incubated overnight at 4°C with 0.2 μg/ml of a goat polyclonal antibody against PCPE-1 (AF2239; Bio-Techne) in PBS, 0.05% Tween-20, 5% skimmed milk. After washing with PBST (PBS plus 0.05 % Tween-20), a secondary rabbit anti-goat antibody coupled to HRP (P0449; Dako) was added for 1 hour (dilution 1:10,000). Finally, after washing, membranes were incubated for 5 min at room temperature with the Amersham ECL Select peroxidase substrate (RPN2235; GE healthcare), and imaged with the Fusion camera. PCPE-1 expression levels were determined using the ImageQuant software (version 8.2, Cytiva) and normalized by total protein amount.

### Immunofluorescence

Paraffin-embedded tissue samples were cut in 4 μm sections. Dewaxing and rehydration were achieved by successive baths in methyl cyclohexane, decreasing ethanol concentration solutions and water. Antigen was retrieved by treatment with a PBS solution of 0.1 % trypsin (T0303, Sigma,) and 0.1 % calcium chloride. After 3 washes, sections were blocked in 4 % BSA (A3059, Sigma) in PBS for 1 h and stained overnight with an anti-PCPE-1 goat polyclonal antibody (12.5 ng/μL, AF2239, Bio-Techne) diluted in 4 % BSA in PBS. After 3 washes and a 1-hour incubation with the secondary anti-goat antibody (3.3 ng/μL, 705605003; Jackson ImmunoResearch) and DAPI (1 μg/μL, 10-50A, Euromedex), sections were washed, mounted in Fluorescence Mounting Medium (F6182, Sigma), dried in the dark and observed using an Eclipse Ti-E fluorescence microscope (Nikon).

### Rat PCPE-1 production and purification

Rat PCPE-1 DNA (NM_019237.1) fused to an N-terminal 8xHis Tag was synthesized by Twist Biosciences. The DNA sequence was then inserted into the pHL-sec vector^37^, between the AgeI and XhoI sites. Rat PCPE-1 was expressed in 293-F cells. First, cells were grown at 37°C, 125 rpm with 8% CO_2_ in FreeStyle^™^ 293 expression medium (Gibco, 12338018) to a cell density of 10^6^ cells/ml. Transfection was performed using linear polyethylenimine (PEI MAX; MW 40,000; Polysciences) and a DNA:PEI ratio of 1:3 (w/w). Three days after transfection, the supernatant was harvested by centrifugation for 20 min at 10,000 g and then loaded onto 5 ml Immobilized Metal-ion Affinity chromatography (IMAC, Ni-NTA Agarose, Qiagen) pre-equilibrated in 20 mM Hepes pH 7.4, 0.3 M NaCl. Following a washing step using the same buffer with 10 mM added imidazole, proteins were eluted with buffer containing 200 mM imidazole. Fractions containing PCPE-1 were then pooled, concentrated with a 10 kDa Amicon ultra-centrifugal filter (Millipore) and purified by size exclusion chromatography on a HiLoad Superdex 75 16/600 (GE healthcare) in 20 mM Hepes pH 7.4, 0.3 M NaCl, 2.5 mM CaCl_2_. Purified rat PCPE-1 concentration (ε= 44350 M^−1^cm^−1^) was determined using a nanodrop ND-2000 (Thermofisher).

### ELISA

A 96-well maxisorp plate (Nunc) was coated overnight at 4°C with a home-made polyclonal rabbit antibody^18^ against PCPE-1 (100 ng per well in 0.2 M carbonate buffer pH 9.2). Wells were then blocked for 1 h at room temperature with 5 % skimmed milk in TBS (50 mM Tris-HCl pH 7.5, 150 mM NaCl). After 3 washes with TBST (TBS plus 0.05 % Tween-20), known amounts of purified rat PCPE-1 or rat plasma samples diluted 10-fold in TBS buffer supplemented with 2.5 mM CaCl2 were added. After incubation for 90 min at room temperature, wells were washed with TBST and rat PCPE-1 was detected using a goat polyclonal anti-PCPE-1 antibody (AF2239; BioTechne; 500 ng/ml). Antibodies bound to PCPE-1 were then detected by incubation with a HRP-conjugated rabbit anti-goat antibody (P0449; Dako; 50 ng/ml). After washing, 100 μl of 3,3’,5,5’-tetramethylbenzidine (TMB) Elisa substrate (T0440; Sigma) was added and the reaction stopped after 10 min using 2 M H_2_SO_4_. The optical density (OD) was measured at 450 nm. PCPE-1 concentration in plasma samples was determined using a second degree polynomial calibration curve.

### Statistics

All statistical analyses were performed using the GraphPad Prism software. Control and fibrotic groups were compared using Student’s t-test (2-tailed). Data were tested for normality with the Shapiro-Wilk test (unless otherwise stated) and for equality of variances with the F-test, and are reported as means ± SD. In the case of unequal variance, Welch correction was performed. p < 0.05 was considered statistically significant.

## (3) Results

### Rat physiological and cardiac function parameters are severely altered by the “AngII + L-NAME” treatment

Rat body weight evolution was similar in the control and “AngII + L-NAME” groups during the first 14 days of treatment. Mean body weight was 303 ± 33 g in the control group vs 297 ± 21 g in the “AngII + L-NAME” group after 2 weeks. After the subcutaneous implantation of angiotensin II-loaded osmotic minipumps, body weight gain was delayed in the “AngII + L-NAME” group as compared to control animals. However, the difference between groups was not significant after 28 days, i.e. at the end of the treatment (Table 2). The mean AngII dose effectively administered was calculated to take into account body weight evolution during the last 2 weeks of the protocol and was 173 ± 3 μg/kg/d.

**Table 2.** Physiological, morphological and cardiac function parameters. Data were tested for normality using Kolmogorov-Smirnof test and are expressed as mean ± SD.

|  | Control | AngII + L-NAME | *p-value* |
| --- | --- | --- | --- |
| **Body weight (g)** | 323 ± 39 | 292 ± 35 | 0.16 |
| **End-diastolic volume**<br>**(mm<sup>3</sup>)** | 377 ± 41 | 287 ± 61 | 0.014 |
| **End-systolic volume**<br>**(mm<sup>3</sup>)** | 114 ± 16 | 137 ± 45 | 0.28 |
| **Ejection fraction (%)** | 69.1 ± 6.9 | 52.5 ± 11.5 | 0.013 |
| **Heart weight (mg)** | 685 ± 111 | 829 ± 36 | 0.008 |
| **Left ventricle volume**<br>**(mm<sup>3</sup>)** | 393 ± 53 | 509 ± 54 | 0.004 |
| **Left ventricle thickness**<br>**(mm)** | 1.48 ± 0.14 | 1.90 ± 0.28 | 0.009 |

Cardiac function evaluated by MRI was modified in the “AngII + L-NAME” group compared to the control group. While the end-systolic volume was only slightly increased, the end-diastolic volume was reduced by more than 20 % in “AngII + L-NAME” treated rats. This resulted in a significant reduction of the ejection fraction (Table 2).

In addition, we observed that the left ventricle volume was significantly elevated in the “AngII + L-NAME” group, as shown by the results obtained by both MR image analyses and tissue weighing at harvesting. As a consequence of diastolic dysfunction and left ventricle volume increase, the left ventricle wall was thicker in the “AngII + L-NAME” group than in the control group (Table 2 and Fig. 1).

**Figure 1.**
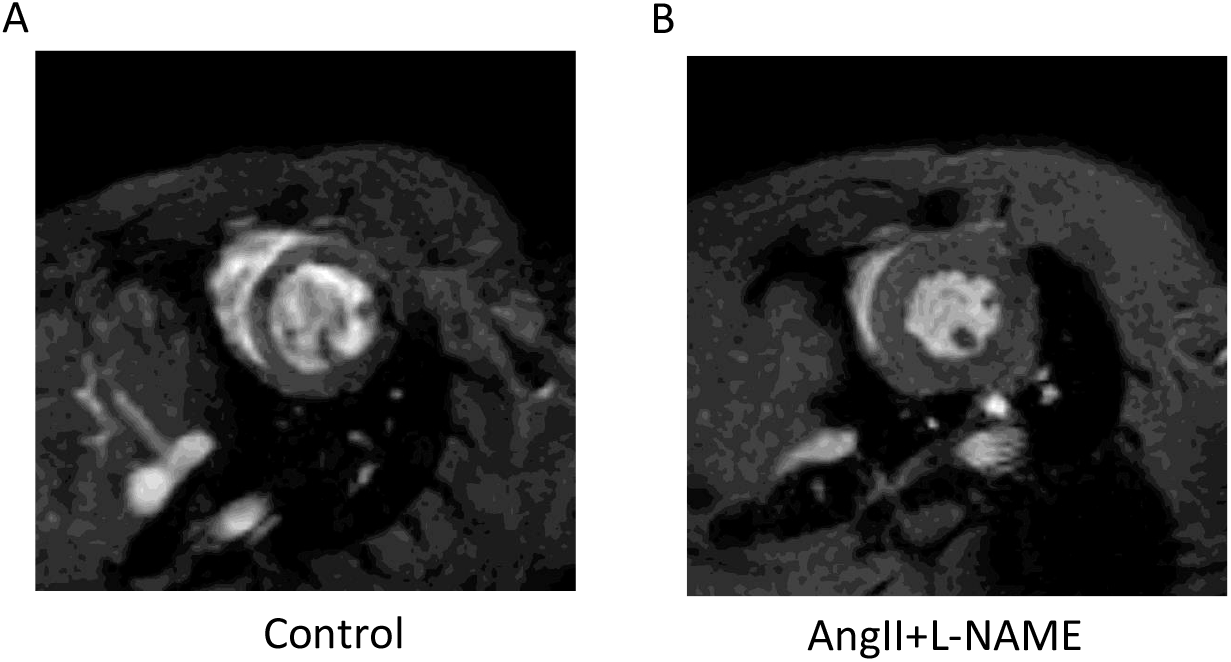
Representative end-diastole MRI images in short axis view obtained in control (A) and “AngII + L-NAME”-treated rats (B).

### Cardiac fibrosis is induced by treatment with AngII and L-NAME

Control animals did not display any sign of fibrosis as evaluated by Masson trichrome or picrosirius red staining (Fig. 2). However, in “AngII + L-NAME” animals, fibrotic regions could be easily delineated using both staining methods (Fig. 2A and 2B). In this group, fibrosis was distributed throughout the ventricular wall and different types of fibrosis could be detected within the same animal cardiac section with 1/ zones typical from reactive interstitial fibrosis with fibrous tissue surrounding muscle cells and 2/ larger patches of fibrotic tissue. Quantification of fibrosis after staining with Masson trichrome or picrosirius red demonstrated the substantial expansion of the fibrotic area in the “AngII + L-NAME” group when compared to the control group (respectively 12.7 ± 6.7 % vs 1.0 ± 0.8 % for Masson trichrome and 10.1 ± 3.6 % vs 3.6 ± 0.7 % for picrosirius red; Fig. 2D and 2E).

**Figure 2.**
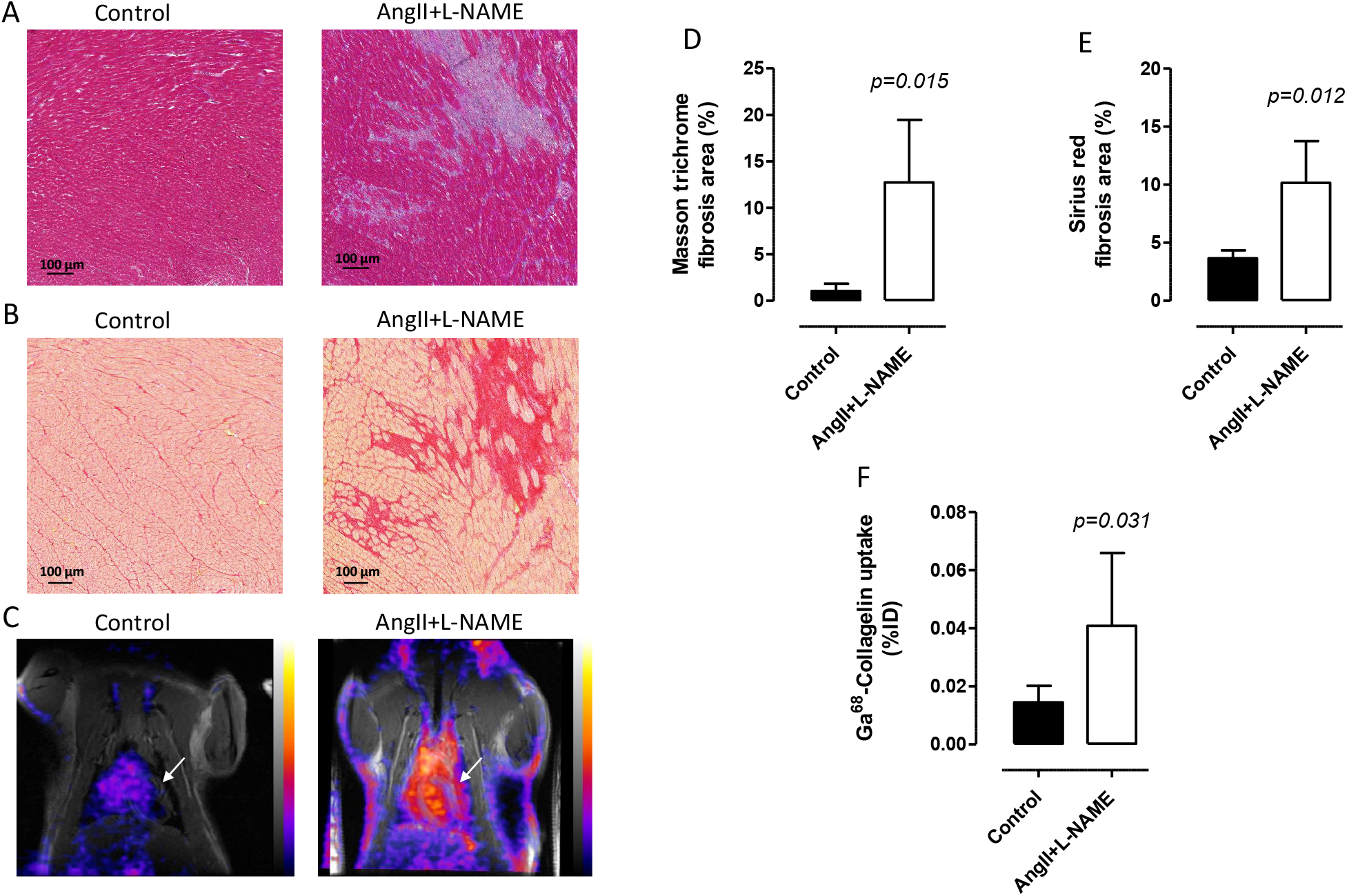
Characterization of left ventricular fibrosis. A – Representative views of Masson Trichrome staining in the left ventricle from control and “AngII + L-NAME” treated rats after 4 weeks of treatment (Magnification x20). B – Representative views of picrosirius red staining obtained as in A. C – Representative PET-MRI co-registered coronal view of a T1-weighed MRI (grey scale) and ^68^Ga-collagelin PET image (color scale) (white arrows indicate heart). D – Fibrosis quantification evaluated in Masson Trichrome-stained left ventricle sections. E – Fibrosis quantification evaluated in picrosirius red-stained left ventricle sections. F – Left ventricle uptake of ^68^Ga-collagelin in control and “AngII + L-NAME” groups measured by radioactivity γ-counting. Data are expressed as mean ± SD.

Finally, we evaluated a PET tracer of collagen, the ^68^Ga-Collagelin^35^. PET images and gamma counting of the cardiac tissue indicated a significant increase of the ^68^Ga-collagelin uptake in the heart of the “AngII + L-NAME” group compared to the control group (Fig. 2C and 2F).

### Genes involved in collagen biosynthesis are up-regulated during cardiac fibrosis

In line with the onset of fibrosis, quantitative gene expression analysis of the rat cardiac samples showed a significant up-regulation of selected genes involved in collagen biosynthesis (Fig. 3). The “AngII + L-NAME” treatment resulted in a 15.9-fold up-regulation of the *Col1a1* gene expression, whereas *Pcolce, Col3a1* and *Bmp1* genes were also significantly increased in fibrotic samples as compared to control samples, by factors of 3.4, 7.8 and 2.8 respectively. On the contrary, the expression of the *Pcolce2* gene encoding the PCPE-2 protein, which shares 41 % identity with PCPE-1 and was previously shown to have a similar enhancing activity at least *in vitro*^38^, was not modified by the treatment.

**Figure 3.**
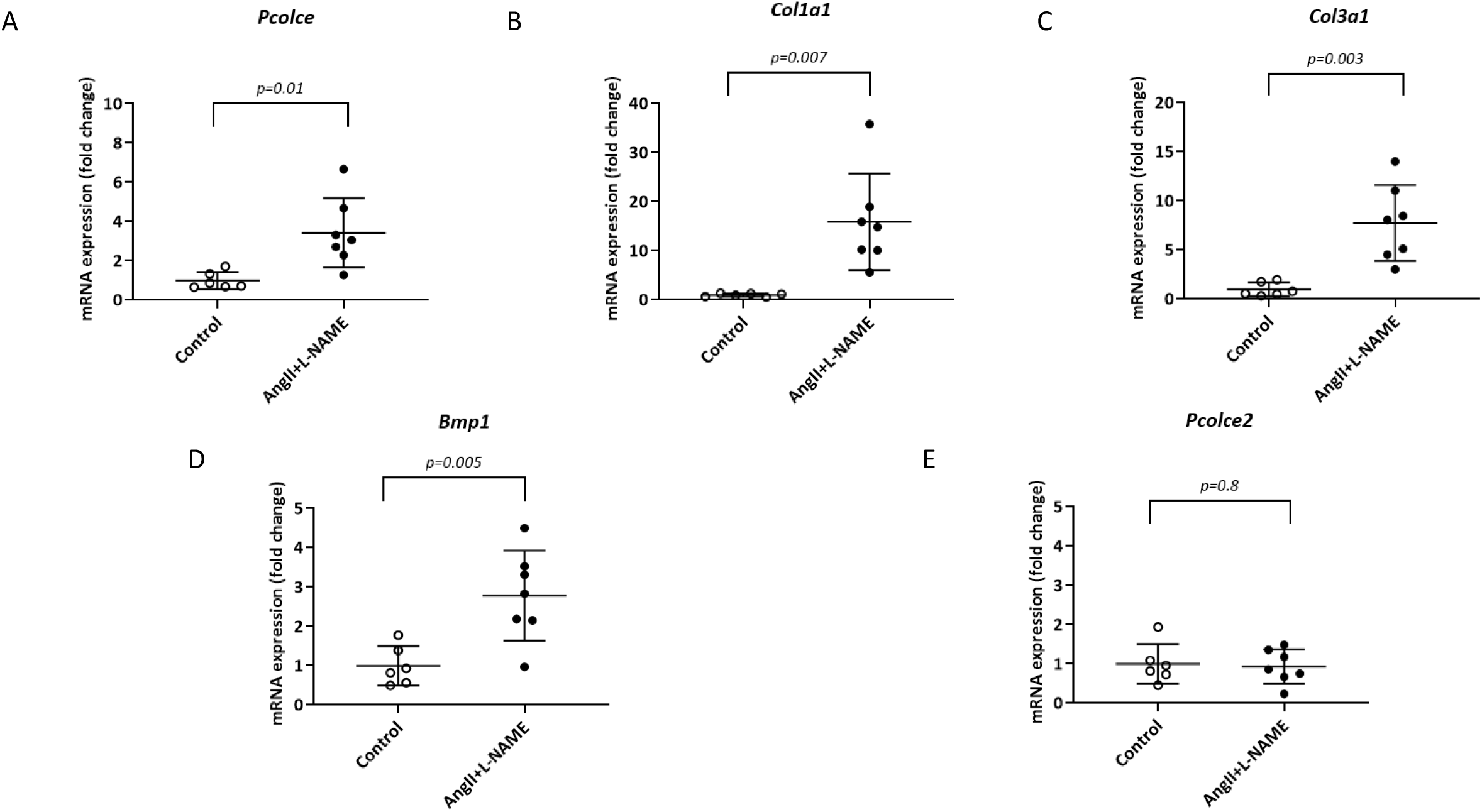
Expression levels of *Pcolce, Col1a1, Col3a1, Bmp1* and *Pcolce2* in the hearts of rats treated or not with AngII + L-NAME (RT-qPCR). The expression levels in the treated rats (n= 7) were compared to the expression levels in the controls (n=6) and analyzed with Student’s t-test with Welch correction (mean ± SD).

### PCPE-1 is increased in fibrotic hearts and correlates with fibrosis

To further explore the role of PCPE-1 in fibrosis, the protein amount was assessed in rat heart tissue using immunoblotting (Fig. 4A). After quantitation and normalization to total protein, PCPE-1 level was found to be 6.1-fold higher in the treated rats compared to controls (P=0.002) (Fig. 4B). We then tested if PCPE-1 levels were directly linked to the level of fibrosis in the rat hearts. The collagen volume fraction (CVF), which is representative of fibrosis levels, varied from 3 % to 15 % based on picrosirius red staining. An excellent correlation was found between PCPE-1 levels and the CVF when all rats, from both control and treated groups, were included (r=0.92, P<0.0001) (Fig 4C).

**Figure 4.**
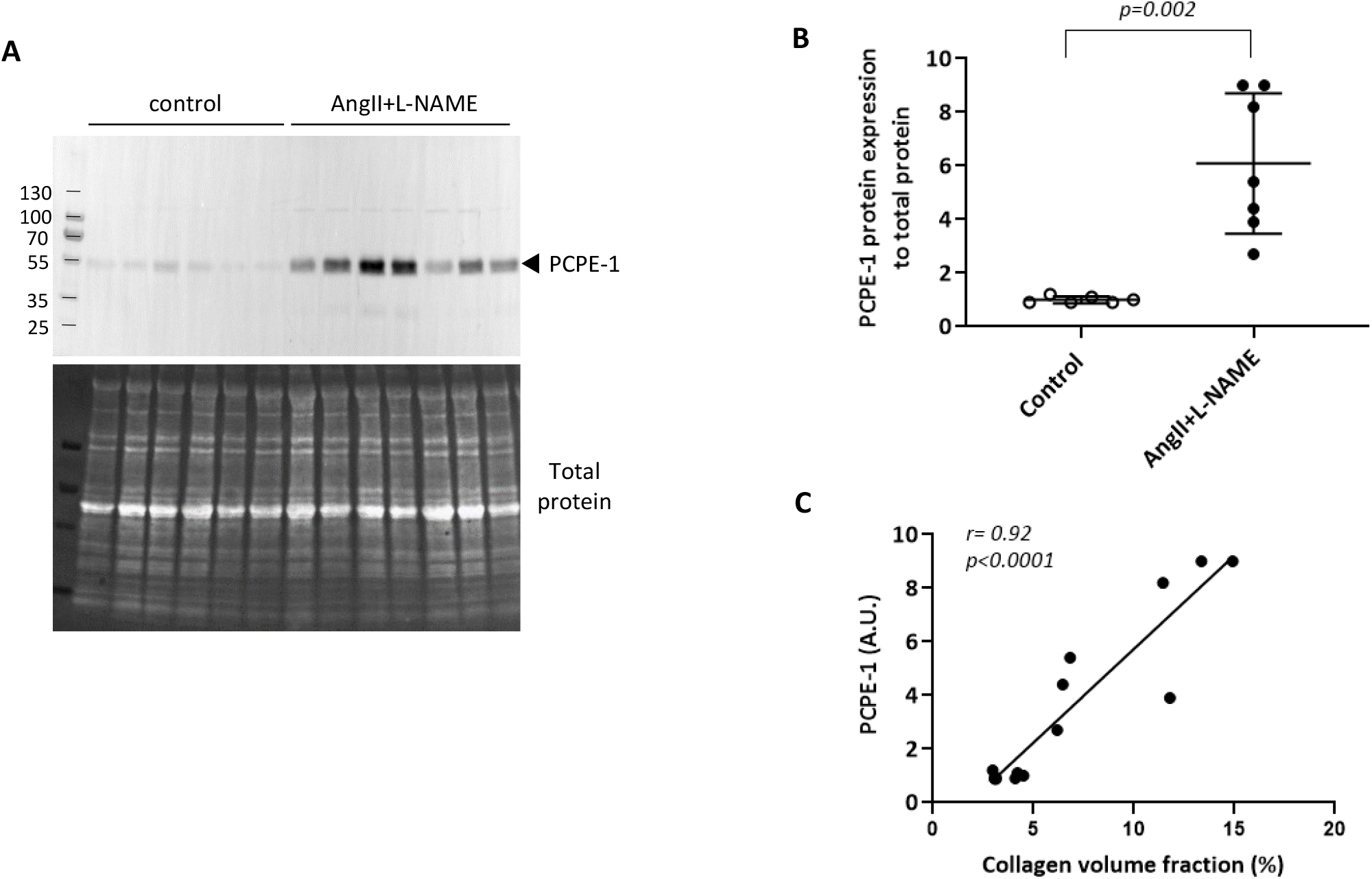
Analysis of PCPE-1 in cardiac tissue. A – Immunoblot showing PCPE-1 content in protein extracts from cardiac tissue in control (C1 to C6) and treated rats (F1 to F7). Lower panel: stain-free imaging of the same membrane. B – Comparison of PCPE-1 level in control and treated rats. Mean ±SD, Student’s t-test with Welch correction. PCPE-1 was quantitated by densitometry and normalized to total protein (labeled by stain free) in 3 separate immunoblots. C – Correlation between PCPE-1 abundance evaluated by immunoblotting and collagen volume fraction (%) determined by picrosirius red staining (linear regression). Each individual dot corresponds to a single rat.

### PCPE-1 localizes to fibrotic areas in the myocardium

PCPE-1 precise location in the myocardium was then evaluated by immunofluorescence. Whereas PCPE-1 was barely visible in control rats, it was easily detected in the “AngII + L-NAME” group (Fig. 5, panel A). Attempts were then made to simultaneously stain collagen I but a high background signal was observed by immunofluorescence and we failed to get coherent data when comparing collagen detection by fluorescence (not shown) and by histological methods (Fig. 2A and B). Interestingly however, the areas which were stained by the anti-PCPE-1 antibody showed a very good overlap in serial sections with the areas stained with picrosirius red (Fig. 5), showing that PCPE-1 is particularly abundant in fibrotic areas of the myocardium.

**Figure 5.**
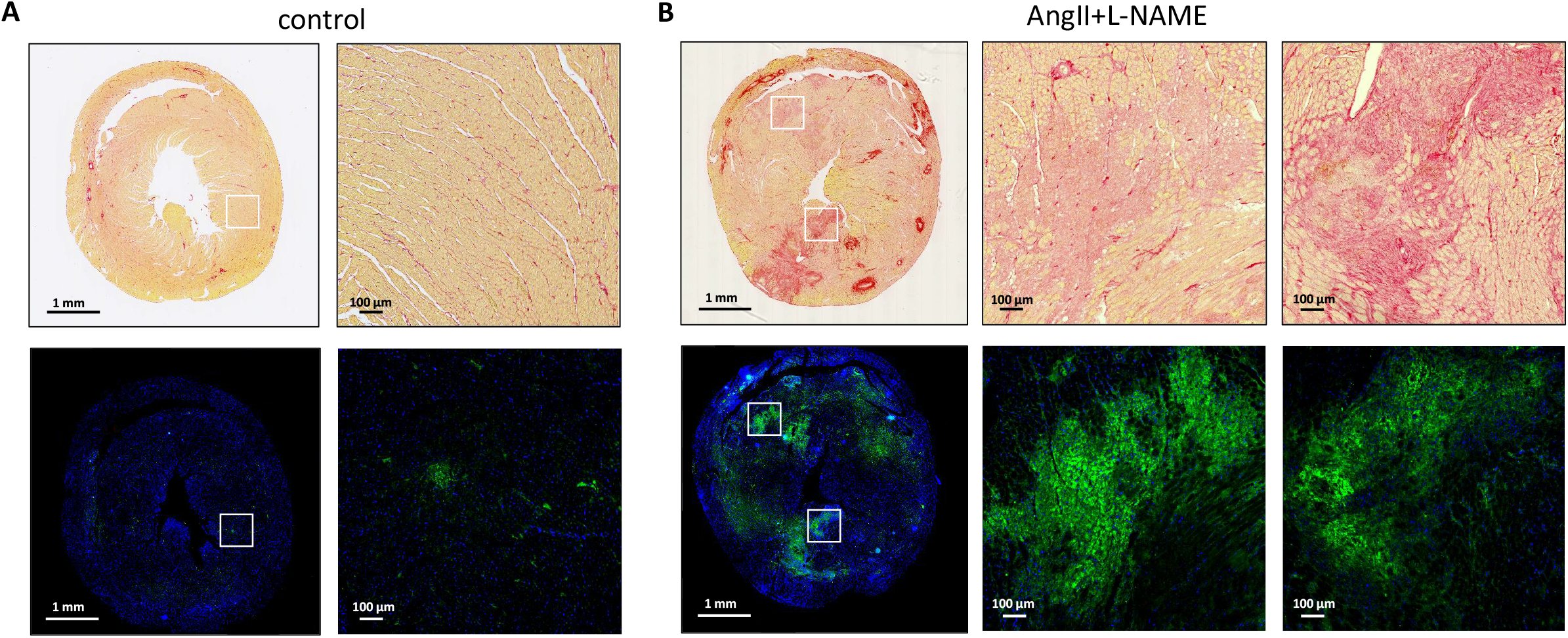
Immunostaining of cardiac sections from control and treated rats with an antibody specific for PCPE-1 and comparison with collagen staining. Collagen was stained with picrosirius red and PCPE-1 (green) was detected in serial sections of the same hearts with Alexa647 conjugated secondary antibody at 680 nm. Nuclei are labeled with DAPI (blue). Representative views from A – control and B – “AngII + L-NAME” treated rats at two different magnifications.

### Plasma levels of PCPE-1 are elevated in fibrotic rats

To measure PCPE-1 concentration in the plasma, a sensitive sandwich ELISA assay was developed (adapted from^25^), using a rabbit polyclonal antibody directed against the NTR domain of PCPE-1 to capture the antigen and a goat polyclonal antibody to the CUB1CUB2 region of PCPE-1 for its detection. A calibration curve, using known amounts of purified recombinant rat PCPE-1, was established, compatible with a 10-fold dilution of plasma samples. The average plasma concentration of PCPE-1 was found to be 201 ± 11 ng/ml in the control rat group (n=6), whereas in the fibrotic group, this concentration was significantly elevated, reaching 228 ± 23 ng/ml (P=0.02, n=7) (Fig 6).

**Figure 6.**
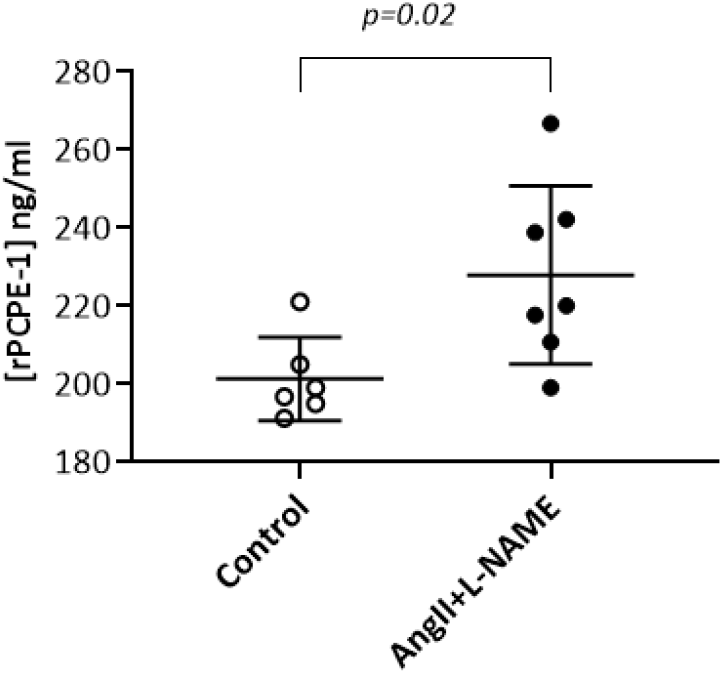
Quantification of circulating PCPE-1 in plasma. Rat PCPE-1 concentration in plasma from control (n=6) and treated rats (n=7) measured by ELISA. Data are representative of two independent measurements, each performed in triplicates. Mean ± SD, Student’s t-test.

## (4) Discussion

Cardiac fibrosis plays a pivotal role in heart failure development and progression. Excessive accumulation of ECM is responsible for fibrosis-driven myocardial stiffening and ECM or ECM-associated proteins are potential fibrosis markers. In the present study, we have characterized a rat model of cardiac fibrosis induced by chronic pressure overload and demonstrated the potential of PCPE-1, a glycoprotein that enhances the processing of fibrillar procollagens, to be used as a biomarker of cardiac fibrosis.

While establishing the animal model, our objective was to reproduce the main characteristics of human cardiac reactive interstitial fibrosis. The latter condition mainly results from increased cardiac overload occurring in the context of aortic stenosis and hypertension and causing collagen accumulation within the cardiac muscle. Therefore, we chose a rat model of pressure overload already described to lead to cardiac fibrosis^34^ which has the advantage of not requiring surgery (lower animal loss) and of being more reproducible than classical models of aortic constriction. With this model, it is reported that if L-NAME administration produces hypertension, the addition of a low dose of AngII exacerbates the remodelling process but does not further increase blood pressure. As compared to the conditions used by Hou and co-workers^34^, we lowered AngII doses (from 225 to 175 μg/kg/d) and extended the treatment for an additional week to reach a total treatment duration of 4 weeks. Reduction and tight control of AngII dose associated with the longer treatment duration led to more reproducible results and to a better evolution of the animals, which otherwise were difficult to maintain until the end of the protocol.

Our experimental model displayed the main features of cardiac interstitial fibrosis that can be observed in the human pathology in which it is commonly associated with left ventricle thickening and diastolic dysfunction with either preserved or reduced ejection fraction^2,3^ (Table 1, Fig. 1). Here, we observed both ventricular thickening and diastolic dysfunction associated with a decrease of ejection fraction which remained above 50 %. More time points would be needed to determine if the decrease of diastolic function precedes ejection fraction reduction, as is often observed^39^. Cardiac dysfunction was also associated with a marked increase of the collagen volume fraction, indicative of a massive deposition of collagens in the treated rats (Fig. 2, 5). Thanks to these characteristics, our improved protocol appears to be a suitable model for the future development and testing of pharmacological compounds targeting myocardial fibrosis^2^.

Consistent with the increased ECM deposition detected by Masson trichrome and picrosirius red staining in the rat samples, we found an up-regulation of the fibrillar collagen genes *Col3a1* (as already noted by Hou *et al*.^34^) and *Col1a1*, accompanied by an elevation of *Pcolce* and *Bmp1* which are both involved in procollagen processing. Up-regulation of *Bmp1* or *Pcolce* has already been reported in other cardiac fibrosis models (myocardial infarction^40^, transverse-aortic constriction (TAC)^30,32^, genetic models^33^) in association with excess collagen deposition. Conversely, we found no differential gene expression of *Pcolce2* despite some reports showing its high expression in the heart^38,41^. This was somewhat unexpected since Baicu *et al*.^41^ described that the PCPE-2 protein (detected by western blotting) is overexpressed in the hearts of mice after TAC and that *Pcolce2* knock-out mice have reduced collagen deposition and myocardial stiffening when subjected to TAC. Human *PCOLCE2* is also reported to be up-regulated in dilated cardiomyopathy^42^, another condition leading to MIF. However, in other fibrotic conditions such as corneal scarring^43^ or liver fibrosis^44^ and contrary to *PCOLCE*, *PCOLCE2* is rather down-regulated or unchanged. These apparent contradictions may reflect different pathological situations and warrant further exploration.

Our gene expression results also demonstrate that collagen synthesis is very active in our model and it is logical to find that *Pcolce*, a gene directly involved in collagen maturation, is up-regulated in this context. The abundance of PCPE-1 in the fibrotic ECM observed by immuno-fluorescence also probably reflects a high level of synthesis by surrounding cells. Interestingly, *Pcolce* was previously shown to be the most differentially expressed fibrogenesis marker among a panel of 67 genes and to allow the early detection of fibrogenesis in models of liver and lung fibrosis^25,45,46^. However, it is not yet known if PCPE-1 could also persist in the extracellular environment after fibrosis reaches its maximum and/or fails to resolve to become installed fibrosis. Several interactions have been described with possible components of the fibrotic ECM (e.g. fibronectin, collagen VI or thrombospondin-1)^47^ that could stabilize PCPE-1 to increase its half-life but it is not clear if PCPE-1 can interact with mature fibrillar collagens. PCPE-1 was shown to co-precipitate with collagen I fibrils^38^ but does not seem to interact with collagen III when the latter is adsorbed in a microtiter plate for ELISA^48^. Whether PCPE-1 is a specific marker of active fibrogenesis or could also reveal established and more stable fibrosis is therefore another important question that needs to be addressed.

Efficient treatment of a disease relies on the availability of effective diagnostic tools that can be used to identify, evaluate and monitor its evolution. Despite its limitations (invasiveness, sampling error…), the analysis of endomyocardial biopsies remains the gold standard to diagnose diffuse cardiac fibrosis^2,9^, and non-invasive procedures to monitor the disease such as immunoassays on body fluids or imaging techniques are critically needed. Contrast-enhanced magnetic resonance (MR) imaging with late gadolinium enhancement and T1 mapping has also been used in recent years for fibrosis evaluation by imaging^2,9,49^. However, it is not fully satisfying since gadolinium locates into the extracellular space (filling effect) and indirectly detects fibrotic regions. Moreover, if MRI is used in clinical practice for the detection of myocardial infarction lesions (localized replacement fibrosis), it is not efficient to detect diffuse interstitial fibrosis. Recently, more specific molecular imaging tracers directly targeting fibrosis have been reported^14^. Here we have used ^68^Ga-Collagelin^14,35,49^ to probe collagen deposition. Collagelin uptake has already been shown to specifically occur in areas of histologically-confirmed fibrosis in various models including myocardial infarction^14,35,49,50^. Our study confirms that the *in vivo* targeting of collagen may be a reliable tool to detect fibrotic lesions in the heart and highlights the potential of ^68^Ga-collagelin PET imaging. Nevertheless, this approach also suffers from some limitations. First, the access of radiotracers to areas of dense and highly cross-linked collagen patches may be impaired by their poor vascularisation, thus explaining the relatively low *in vivo* uptake of ^68^Ga-collagelin in fibrotic hearts compared to controls (e.g. less than 0.05% of the injected dose in Fig. 2). Second, targeting collagen accumulation may not be the optimal objective as massive collagen deposition is a late event in fibrogenesis which might reflect a stage of the disease already too advanced for therapeutic intervention and efficacy. A better option would be to focus on early players of the fibrotic process such as PCPE-1.

In this context, there is still a clear potential for the identification of novel circulating biomarkers of cardiac fibrosis. Among all the proteins that have been suggested so far, only the C-propeptide of collagen I (PICP) and the N-propeptide of collagen III (PIIIPN) have been clearly correlated with histological results and are used in (pre)clinical studies, but even those are not completely specific of cardiac fibrosis and are not used in clinical practice^2,12^. Also, it is highly probable that a combination of biomarkers will be needed to properly monitor the evolution of the disease and PCPE-1 could be one of these mechanistically-driven biomarkers integrated into those combinations. Indeed, the over-expression of PCPE-1 in fibrotic diseases^27,43,51^, including cardiac pathologies^29^, is already well documented, and PCPE-1 has been previously suggested as a biomarker of cardiac fibrosis^28,29,52,53^. Similar to what is described for other fibrotic conditions, including both patient samples^51,54^ and animal models^25,45^, and for the first time in the context of heart fibrosis, we have found an increased concentration of circulating PCPE-1 in the fibrotic rats compared to controls. Average plasma levels (around 200 ng/ml) detected in normal rats were very similar to those reported for mice^25^ or human^51,54^, but much higher than those previously reported by Ippolito *et al*.^45^ (1.5 ng/ml) in rats. Whereas the latter study used a commercial Elisa kit, our ELISA assay was developed using well-characterized antibodies, and above all, freshly prepared rat PCPE-1 expressed in mammalian cells as a standard. This probably accounts for the observed discrepancy and stresses the importance of well-defined protein and antibody sources when devising a biomarker assay but, most importantly, our results also show that PCPE-1 is detected and elevated in the plasma of rats undergoing myocardial fibrosis.

Histological validation and correlation with CVF is a prerequisite to biomarker validation, which should reflect the extent of fibrosis and the progression of the disease^12^. Here we show that not only is PCPE-1 expression increased in the heart of the fibrotic rats compared to untreated rats, but also that PCPE-1 levels (detected by western blotting) are directly and strongly correlated to fibrosis levels (measured by picrosirius red, Fig. 5). A similar result was previously obtained in a murine model of replacement fibrosis^30^ and suggests that, despite the different models, common players, including PCPE-1, are involved in excessive collagen deposition. Further, analysis of biopsies by immunofluorescence confirmed that PCPE-1 mainly localizes to fibrotic areas, including patterns of interstitial fibrosis. These results show that PCPE-1 holds the characteristics needed to define a circulating biomarker of cardiac fibrosis^12^. Whether this can be translated to human pathology remains to be established, but the added value of using PCPE-1 in a combination including the validated biomarkers derived from collagen turn-over (PICP, PIIINP) deserves further investigation.

## (5) Acknowledgments

Paraffin-embedding, Masson trichrome and picrosirius red staining of cardiac sections were performed by the CellImaP core facility, INSERM U1231 (Dijon, France). We acknowledge the contribution of the AniRA Genetic Analysis facility from the SFR Biosciences (UMS3444/CNRS, US8/Inserm, ENS de Lyon, UCBL) for the access to the Rotor-Gene equipment and initial training on how to perform qRT-PCR. We also would like to thank Leah Berger and Laeticia Moisy for their help with the preliminary optimizations of PCPE-1 detection.

## Data availability statement

The data underlying this article will be shared on reasonable request to the corresponding authors.

